## Supplementary materials for "A multiscale computational framework for the development of spines in molluscan shells"

### Supplementary Material - A multiscale computational framework for the development of spines in molluskan shells

#### 1 Biochemical model

The biochemical model we use is taken from Fowler et al. [1], Equation (8) in the main text. In dimensional form, this reads as:

$$\begin{aligned}\frac{\partial A}{\partial T} &= \rho S\left(\frac{A^2}{1+\kappa A^2}+\rho_0\right)-\mu A + D_A\frac{\partial^2 A}{\partial X^2}\\ \frac{\partial S}{\partial T} &= \sigma-\rho S\left(\frac{A^2}{1+\kappa A^2}+\rho_0\right)-\nu S + D_S\frac{\partial^2 S}{\partial X^2}.\end{aligned} \tag{1}$$

Here, $A(X,T)$ is the concentration of the activator, a function of the spatial variable, $X$, and time, $T$, while $S(X,T)$ is the concentration of the substrate. The activator diffuses with diffusion constant $D_A$ and decays at rate $\mu$, while the substrate diffuses with rate $D_S$, decays at rate $\nu$ and is produced at constant rate $\sigma$. The term

$$\rho S\left(\frac{A^2}{1+\kappa A^2}+\rho_0\right)$$

describes an autocatalytic process for production of activator. This requires the presence of substrate, has a baseline production rate, $\rho_0$, and has an additional production depending nonlinearly on the concentration of activator: quadratically for small values of $A$ and saturating at large activator concentrations, with saturation depending on the parameter $\kappa$. We scale the variables as:

$$A=\left(\frac{\sigma}{\rho}\right)^{2/3}a,\ \ T=\frac{1}{\rho^{1/3}\sigma^{2/3}}t,\ \ S=\left(\frac{\sigma}{\rho}\right)^{1/3}s, \tag{2}$$

from which we obtain the following nondimensional equations:

$$\begin{aligned}\frac{\partial a}{\partial t} &= \frac{sa^2}{1+c_1a^2}+c_2 s - c_3 a + d_1\frac{\partial^2 a}{\partial x^2}\\ \frac{\partial s}{\partial t} &= 1-\frac{sa^2}{1+c_1a^2}-c_4 s + d_2\frac{\partial^2 s}{\partial x^2},\end{aligned} \tag{3}$$

as presented in the main text. Here the variables $\{c_1, c_2, c_3, c_4, d_1, d_2\}$ are defined by

$$c_1\kappa a_0^2,\ \ c_2 = \frac{\rho\rho_0 s_0\tau}{a_0},\ \ c_3 = \mu\tau,\ \ c_4 = \tau(\rho\rho_0+\nu),\ \ d_1 = \frac{D_A\tau}{L^2},\ \ d_2 = \frac{D_S\tau}{L^2}, \tag{4}$$

where $a_0 = \left(\frac{\sigma}{\rho}\right)^{2/3}$, $s_0 = \left(\frac{\sigma}{\rho}\right)^{1/3}$, $\tau = \frac{1}{\rho s_0 a_0}$, and $L$ is the dimensional length of the domain. These equations are solved with periodic boundary conditions and uniform initial concentrations of unity, plus a small degree of noise added to $a$ to enable pattern-forming when the homogeneous state is unstable.

### 2 Mechanical model and computational approach for spine formation

In this section we first present the full mechanical model of mantle growth and deformation, which is derived from the theory of morphoelastic rods [2, 3], and also describe the computational approach to generate the different *in silico* spine morphologies.

#### 2.1 Mechanical model

The mantle edge is modeled as a growing planar, inextensible elastic rod. The shape of the rod is described by its centreline curve in two-dimensional Cartesian geometry, such that $\mathbf{r} = x\mathbf{e}_x + y\mathbf{e}_y$.

$$\lambda = \alpha\gamma \qquad \iff \qquad \frac{\partial s}{\partial S_0} = \frac{\partial s}{\partial S}\frac{\partial S}{\partial S_0}, \tag{5}$$

where $\lambda$ is the total stretch of the rod; $\alpha$ is the elastic stretch; and $\gamma$ is the growth stretch. For an inextensible rod, the elastic stretch $\alpha = \frac{\partial s}{\partial S} \equiv 1$. Therefore, in this case, the total stretch, $\lambda$, is equal to the growth stretch $\gamma$:

$$\alpha = \frac{\partial s}{\partial S} = 1 \qquad \implies \qquad \lambda = \gamma(S_0, t) = \frac{\partial s}{\partial S_0}. \tag{6}$$

We model the incremental growth of the mantle edge as a Gaussian function:

$$\frac{\dot\gamma}{\gamma} =: g(s) = g_1 \exp\left(-\frac{(s-\frac{l}{2})^2}{2g_2^2}\right), \tag{7}$$

where the overdot represents partial differentiation with respect to $t$.

In order to describe the resistive effects of the shell edge, we model the shell as a smooth curve, $\mathbf{p} = p_x\mathbf{e}_x + p_y\mathbf{e}_y$, which is tethered to the growing rod. We assume that a linear elastic constitutive relation for shell attachment to the mantle and assume that additional shell material is secreted and calcified over time due to growth of the mantle. Note that we assume that there is no twisting effect induced by $\mathbf{p}$; hence, $p_z \equiv 0$. Let $\mathbf{n} = n_x\mathbf{e}_x + n_y\mathbf{e}_y$ and $\mathbf{m} = m\mathbf{e}_z$ denote, respectively, the resultant force and bending moment in the growing rod. Here, the primes $'$ denote (partial) differentiation with respect to the initial arc length parameter $S_0$ (hence the presence of factors of $\gamma$ in Equations (10)–(15)). The bending moment, $m$, is assumed to depend on curvature via the constitutive relation:

$$m = EI\gamma^{-1}\theta' = \frac{Ewh^3}{12\gamma}\theta', \tag{8}$$

where $E$ is the mantle stiffness and $I$ is the second moment of area, where we have assumed a rectangular cross-section.

We adopt the assumption that the mantle is not explicitly growing (increasing in mass e.g. via cell division), but rather stretching in the direction along the mantle edge; conservation of volume then implies that the thickness must decrease proportionally, and consequently the bending stiffness will decrease as well, as it is proportional to cube of the thickness for a rectangular cross-section. This assumption comes from the observation that the length of the mantle margin just after spine production is approximately the same as before. Therefore, if the mantle were in fact growing during spine production, there would have to be significant resorption (negative growth) at the end of the spine formation process, which seems unlikely. We note that the alternative assumption, that the mantle does in fact grow, would also be feasible to analyse within our modelling framework. However, in such a case, growth and stiffness would not naturally be linked, and one would have to decide how to separately determine their profiles, perhaps requiring further species in the biochemical component.

The above implies that during spine formation, the total mantle tissue material is conserved; that is, any growth, $\gamma$, is balanced by a contraction in the cross-sectional height, $h$. This implies that $h(t) = h\gamma^{-3}$.

The model is scaled using the following nondimensionalization [2]:

$$\begin{aligned}(S_0^*, x^*, y^*, p_x^*, p_y^*) &= L_0(S_0, x, y, \Delta, p_x, p_y)\\ (n_x^*, n_y^*) &= E(I/L_0^2)(n_x, n_y),\\ \sigma^* &= E(I/L_0^3)\sigma,\\ m^* &= E(I/L_0)m,\\ t^* &= \tau t.\end{aligned} \tag{9}$$

Here, $L_0$ is the initial rod length; $E$ is the Young's Modulus; $I = wh^3/12$ is the (second) moment of inertia, for an assumed rectangular cross-section with width $w$ and height $h$; and $\tau$ is a typical timescale for tissue growth, on the order of a day (i.e. $\tau = 24$ hours). Therefore, the full set of equations describing geometric constraints and balance of linear and angular momentum are:

$$x' = \gamma \cos\theta, \tag{10}$$

$$y' = \gamma \sin\theta, \tag{11}$$

$$n_x' = \gamma k(x - p_x), \qquad \dot{p}_x = \eta(x - p_x), \tag{12}$$

$$n_y' = \gamma k(y - p_y), \qquad \dot{p}_y = \eta(y - p_y), \tag{13}$$

$$\theta' = \gamma^4 m, \tag{14}$$

$$m' = \gamma(n_x \sin\theta - n_y \cos\theta). \tag{15}$$

where the nondimensional parameter, $k = 12k_f L_0^4/(wh^3)$, describes the strength of resistance by the shell edge.

To close the system, we assume the rod is clamped horizontally at the boundaries:

$$x(0) = 0, \quad y(0) = 0, \quad \theta(0) = 0, \qquad x(1) = 1, \quad y(1) = 0, \quad \theta(1) = 0. \tag{16}$$

#### 2.2 Computational simulation details

For all simulations, the shell stiffness scaling parameter $k_f$ has been set to $k_f = 0.01$, which produces a mode 1 shape upon buckling. This shape represents the formation of a single spine. Following previous works [2], we assume an initial rectangular cross-section of $h = 0.7mm$ and width, $w = 0.7mm$, and an initial rod length, $L_0^* = 1cm$.

Spine formation is triggered through growth-induced buckling. As we assume the rod is inextensible, buckling occurs for any time $t > 0$ for mantle growth such that the total rod length, $l > 1$. All time derivatives are discretised using the Forward Euler method, with a timestep $\delta t$. To ensure that there is no jump in solution branch and stability during spine simulation, we must set the time step increment, $\delta t$, to be sufficiently small. We found that setting $\delta t$ in the range $\delta t = 0.01 - 0.05$ was sufficient. As such, we simulated spine formation over times $t_i = k\delta t$, $k = 0, \ldots, N$, where $N$ is defined such that at time $t_{N+1}$, self-contact of the mantle edge occurred. At each time step, we first update the evolution equation for mantle growth, Equation (7). Assuming quasi-static mechanical equilibrium, the resulting boundary value problem governed by Equations (10)–(15) are solved numerically using the MATLAB package `bvp4c`, which employs a fourth-order spatial collocation method [4]. After obtaining the numerical solution for the mantle edge, $(x(S_0), y(S_0))$, we update the shell edge, $(p_x(S_0), p_y(S_0))$, using the discretised form of Equations (12)–(13). As we are solving a boundary value problem at each time step, we must provide a suitable initial "guess" for the numerical solution. At $t = 0$, when no growth has occurred, the mantle and shell edge are flat, i.e. $x(S_0) = p_x(S_0) = S_0$, $y = p_y = 0$. Therefore, to obtain the first buckled profile, we can use numerical solutions of the the model corresponding to the special case of constant growth, $\gamma(S_0, t) = \gamma > 1$, and a static foundation, $\eta = 0$, as the initial guess. We used the numerical continuation package `AUTO-07p` [5] to generate a guess for the initial solution, where continuation was performed with respect to $\gamma$. For all subsequent times, we used the solution at time $t_k$ as the initial guess for the solution at time $t_{k+1}$.

To speed up simulations, noting that we only need to simulate even solutions of the mantle and shell, we translate the domain of growth to $S_0 \in [-1/2, 1/2]$ and only consider rod growth on the right half-interval, $S_0 \in [0, 1/2]$. As the rod is no longer clamped at $S_0 = 0$, the boundary condition $y(0) = 0$ in (16) no longer applies. However, noting that the $y(S_0)$ and $p_y(S_0)$ must be even about $S_0 = 0$ and the local maximum of the mantle occurs at $S_0 = 0$, it follows that $n_y(0) = 0$ and $\theta(0) = 0$. Therefore, the original boundary conditions (16) are replaced by:

$$x(0) = 0, \quad n_y(0) = 0, \quad \theta(0) = 0, \qquad x\left(\frac{1}{2}\right) = 1, \quad y\left(\frac{1}{2}\right) = 0, \quad \theta\left(\frac{1}{2}\right) = 0. \tag{17}$$

### 3 Linking biochemical and biomechanical models

To link between the two modelling scales, we relate the steady state activator profile, $a_\infty(x) = \lim_{t\to\infty} a(x, t)$, to the incremental growth profile, $g(s)$, as modeled in Equation (7). Here, we note that under periodic boundary conditions, as we have used, the biochemical system may output a profile that is well-approximated by a Gaussian, but shifted up or down by a constant value, $a_c$, say. We neglect this constant value in passing to the biomechanical model, which reflects the notion that it is spatial gradients in concentrations that should produce a non-uniform pattern, and we also apply, if necessary, a horizontal shift so that the profile is centered in the horizontal direction. We thus extract from the steady state profile a best-fit Gaussian, i.e., we find the values $\{a_1, a_2\}$ that best approximate the shifted activator profile, $a_\infty(x) - a_c$, as a Gaussian; that is, we approximate

$$a_\infty(x) = a_c + a_1 \exp\left(-\frac{(x - L/2)^2}{2a_2^2}\right).$$

The best-fit values of $a_1$ and $a_2$ were obtained using the SciPy [6] function Curve$_{\text{fit}}$. From here, we equate $g_2 = a_2$, since both systems are solved in nondimensional form on the same domain length, but we allow for scaling between $g_1$ and $a_1$, taking $g_1 = \alpha a_1$, where $\alpha$ is taken to be the same fixed value for all simulations.

The biochemical model, even in nondimensional form, forms a six-dimensional parameter space $\{c_1, c_2, c_3, c_4, d_1, d_2\}$. Only a subset of this space will form a pattern at all, and only a small subset of that space will form a pattern that can be reasonably approximated as a single Gaussian. In order to maintain computational tractability, we therefore perform two steps prior to running the Quality-Diversity algorithm: (i) we fix the diffusion constants $\{d_1, d_2\}$, and (ii) we define fixed ranges for the other 4 parameters $\{c_1, c_2, c_3, c_4\}$. The diffusion constants are set as $d_1 = 0.0018, d_2 = 0.707$; these values are similar to those given in the paper [1] that originally presented this Activator-Substrate system. A biological rationale for fixing these is that since these describe physical properties of chemicals, they are unlikely to vary between individuals, regardless of environmental conditions. In order to define ranges for the $\{c_i\}$ parameters, we performed cursory simulations of the system (18) within the `Manipulate` command in Mathematica [7], which enables us to vary the parameters within a GUI environment while solving the system up to a large time and monitoring the output profiles. In this way, we defined a range of values for which the system produced profiles that had the desired characteristics.

### 4 Stability boundary of biochemical model

In order to better understand the presence of large white regions in the $\{c_i\}$ parameter space (Fig 4(b)) for which the Quality-Diversity algorithm failed to find a suitable Gaussian profile, we conduct here a standard linear stability analysis of the system (18) in order to identify boundaries in parameter space separating stable regions (for which a pattern will not form) from unstable, pattern-forming regions. The linear stability analysis that follows is very standard (see e.g. [8], Sec 2.3), and is therefore only briefly outlined here. We first express the system in the form

$$\begin{aligned} \frac{\partial a}{\partial t} &= f(a,s) + d_1 \frac{\partial^2 a}{\partial x^2} \\ \frac{\partial s}{\partial t} &= g(a,s) + d_2 \frac{\partial^2 s}{\partial x^2}, \end{aligned} \tag{18}$$

where

$$f(a,s) := \frac{sa^2}{1 + c_1 a^2} + c_2 s - c_3 a,$$

$$g(a,s) := 1 - \frac{sa^2}{1 + c_1 a^2} - c_4 s.$$

The conditions for a diffusion-driven instability come from perturbing about a homogeneous steady state $a = a^*, b = b^*$ satisfying $f(a^*, b^*) = 0 = g(a^*, b^*)$, and requiring that the steady state is stable in the absence of diffusion, but unstable in the presence of diffusion. We define the terms:

$$\mathcal{A} = d_2 f_a + d_1 g_s, \quad \mathcal{B} = f_a g_s - f_s g_a,$$

where subscript denotes partial differentiation, and $\mathcal{A}$, $\mathcal{B}$ are evaluated at the point $(a^*, b^*)$. The condition for stability is given by:

$$\mathcal{A}^2 - 4 d_1 d_2 \mathcal{B} > 0$$

and either $\mathcal{B} < 0$ or $\mathcal{A} > 0$. Points at which these conditions do not hold are linearly unstable, and therefore pattern-forming.

To compare the stability boundary against the computed values from the Quality-Diversity algorithm, we fix $c_4 = 0.005$, as this parameter has been shown to have little to no effect on the profile, vary $c_3$ over the range 0.4 to 1.5, and then compute the stability boundary as a curve in the $c_1$-$c_2$ plane for each value of $c_3$. These curves are plotted in Fig 1, such that points below (above) the curve form the unstable (stable) region. We also include in this plot the full set of $\{c_1, c_2\}$ values produced by the Quality-Diversity algorithm, colored by corresponding $g_1$ value. The envelope of the stability boundary curves is very closely aligned with the boundary of the set of $\{c_1, c_2\}$ values detected from the Quality-Diversity algorithm, suggesting that indeed the white space primarily corresponds to regions in which the system does not create a pattern.

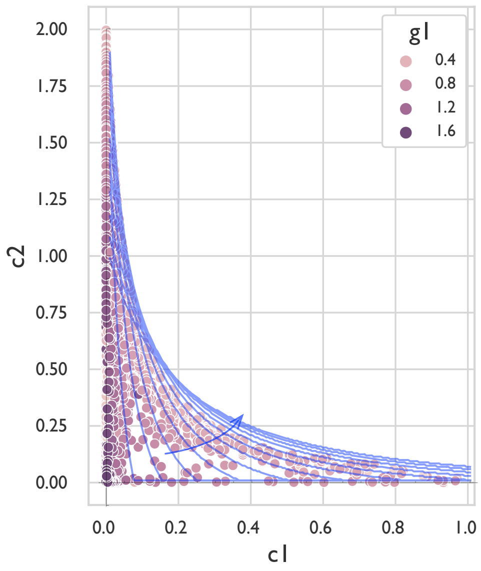


**1.** The full set of $\{c_1, c_2\}$ values produced by the Quality-Diversity algorithm, colored by corresponding $g_1$ value. The blue curves are stability boundary curves for fixed $c_4 = 0.005$ and varying $c_3$, increasing from 0.4 to 1.5 following the arrow.

#### 5 Spine images and extracted values

In this section we report the results of our morphometric analysis of spine shape. We analyzed $N$ Spine images from each location, with $N = 23$ for Hayama, $N = 28$ for Jyogashima, and $N = 16$ for Tateyama. As described in the main text, each image was imported into Mathematica and output from the mechanical model was overlaid and compared, after appropriate scaling, rotation, and translation. Figure 2 shows the spines from Hayama, with a single model output curve overlaid; corresponding plots for Jyogashima and Tateyama are provided in Figures 3 and 4, respectively. Extracted $(g_1, g_2)$ parameters are provided in Tables 1-3. In order to record spine location on the shell edge, as well as spines from the same specimen, we have labelled the spines both by specimen number and location on the shell, such that for a given shell number, spines on the shell edge are labelled A-D from left to right. (In one instance, we have used a spine secreted at an earlier time, thus appearing recessed from the shell edge. This was labelled Hayama 4A(ii) to distinguish from the spine Hayama 4A, which appeared in the same relative location but at the shell edge.)

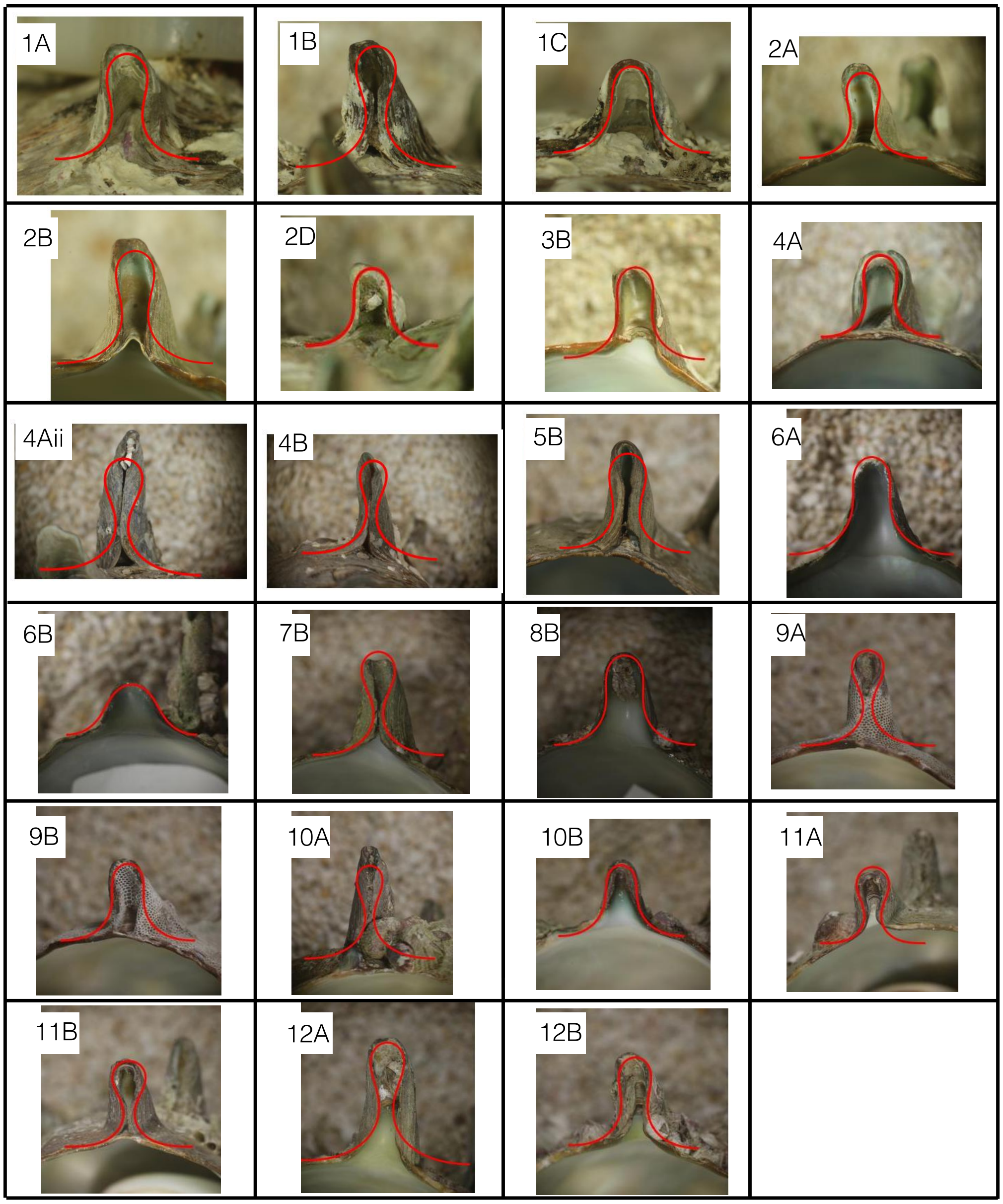


**2.** Spines analysed from Hayama, including a single curve from the tissue model overlaid. Extracted $(g_1, g_2)$ values are given in Table 1.

| Shell | Spine | $g_1$ | $g_2$ |
|---|---|---|---|
| 1 | A | 0.1 | 0.3 |
| 1 | B | 0.1 | 0.25 |
| 1 | C | 0.1 | 0.2 |
| 2 | A | 0.25 | 0.25 |
| 2 | B | 0.1 | 0.25 |
| 2 | D | 0.25 | 0.25 |
| 3 | B | 0.1 | 0.25 |
| 4 | A | 0.1 | 0.3 |
| 4 | A(ii) | 0.1 | 0.2 |
| 4 | B | 0.25 | 0.2 |
| 5 | B | 0.1 | 0.3 |
| 6 | A | 0.1 | 0.2 |
| 6 | B | 0.1 | 0.25 |
| 7 | B | 0.1 | 0.25 |
| 8 | B | 0.1 | 0.25 |
| 9 | A | 0.25 | 0.2 |
| 9 | B | 0.25 | 0.2 |
| 10 | A | 0.1 | 0.2 |
| 10 | B | 0.1 | 0.2 |
| 11 | A | 0.5 | 0.2 |
| 11 | B | 0.25 | 0.3 |
| 12 | A | 0.1 | 0.2 |
| 12 | B | 0.1 | 0.25 |

**Table 1.** Extracted $(g_1, g_2)$ values for the spines from Hayama, corresponding to the images in Fig 2.

### 6 Statistical testing of inferred spine growth rate parameters

We performed statistical testing to infer whether the incremental growth rate parameters, $g_1$ and $g_2$, for each of the Hayama (sample size $N_H = 24$), Jyogashima (sample size $N_J = 28$), and Tateyama (sample size $N_T = 15$). prefectures were distinct. For each pair of sampled prefectures and for the inferred values of $g_1$ ($g_2$), we performed a two-sided Wilcoxon rank-sum test, under the null hypothesis assumption that the distributions of $g_1$ ($g_2$) were equal. For all tests, we assumed a significance level of $p = 0.05$.

The distribution of $g_1$ values from the Jyogashima region were inferred to be significantly higher than those from the Hayama region (test statistic $W = 111.5$, $p-$value $p = 1.202 \times 10^{-5}$), with an estimated difference in $g_1$ of 0.25 (95% confidence interval = $[0.15, 0.40]$). However, the differences in $g_2$ values were estimated to be insignificant ($W = 410, p = 0.1422$), with an estimated difference of $-5.05 \times 10^{-5}$ (95% CI = $[-0.05, 5.52 \times 10^{-5}]$).

We failed to reject the hypothesis that $g_1$ values from the Tateyama region are significantly different from those from the Hayama region (test statistic $W = 214.5$, $p-$value $p = 0.224$), with an estimated difference in $g_1$ of $-6.04 \times 10^{-5}$ (95% confidence interval = $[-2.66 \times 10^{-5}, 3.34 \times 10^{-5}]$). However, the differences in $g_2$ values were estimated to be significant but only marginally smaller from the Tateyama region ($W = 297.5, p = 0.0002$), with an estimated difference of $-0.05$ (95% CI = $[5.52 \times 10^{-5}, 0.05]$).

The $g_1$ values from the Jyogshima region were estimated to be significantly higher

| Shell | Spine | $g_1$ | $g_2$ |
|---|---|---|---|
| 1 | A | 1.0 | 0.2 |
| 1 | B | 1.0 | 0.25 |
| 2 | A | 0.5 | 0.25 |
| 2 | B | 0.5 | 0.25 |
| 3 | A | 0.5 | 0.2 |
| 3 | B | 0.5 | 0.2 |
| 4 | A | 0.5 | 0.2 |
| 5 | A | 0.5 | 0.2 |
| 5 | B | 0.5 | 0.2 |
| 6 | A | 0.5 | 0.3 |
| 6 | B | 0.5 | 0.25 |
| 7 | A | 0.5 | 0.2 |
| 7 | B | 0.5 | 0.2 |
| 8 | A | 0.5 | 0.25 |
| 8 | B | 0.5 | 0.3 |
| 9 | B | 0.5 | 0.25 |
| 10 | A | 0.5 | 0.25 |
| 10 | B | 0.5 | 0.2 |
| 11 | A | 0.5 | 0.3 |
| 11 | B | 0.25 | 0.2 |
| 12 | A | 0.25 | 0.2 |
| 12 | B | 0.25 | 0.2 |
| 13 | A | 0.25 | 0.25 |
| 14 | A | 0.1 | 0.2 |
| 14 | B | 0.1 | 0.2 |
| 15 | A | 0.5 | 0.2 |
| 15 | B | 0.25 | 0.2 |
| 16 | B | 0.5 | 0.2 |

**Table 2.** Extracted $(g_1, g_2)$ values for the spines from Jyogashima, corresponding to the images in Fig 3.

| Shell | Spine | $g_1$ | $g_2$ |
|---|---|---|---|
| 1 | A | 0.1 | 0.2 |
| 1 | B | 0.1 | 0.2 |
| 2 | A | 0.1 | 0.2 |
| 2 | B | 0.1 | 0.2 |
| 3 | A | 0.25 | 0.2 |
| 3 | B | 0.1 | 0.2 |
| 4 | A | 0.1 | 0.25 |
| 4 | B | 0.1 | 0.2 |
| 5 | A | 0.1 | 0.2 |
| 5 | B | 0.1 | 0.2 |
| 6 | A | 0.1 | 0.2 |
| 6 | B | 0.1 | 0.2 |
| 7 | B | 0.1 | 0.2 |
| 8 | A | 0.1 | 0.2 |
| 9 | A | 0.25 | 0.2 |
| 9 | B | 0.1 | 0.2 |

**Table 3.** Extracted $(g_1, g_2)$ values for the spines from Tateyama, corresponding to the images in Fig 4.

than those from the Tateyama region (test statistic $W = 376.5$, $p-$value $p = 5.70 \times 10^{-6}$), with an estimated difference in $g_1$ of 0.40 (95% confidence interval = $[0.25, 0.40]$). However, the differences in $g_2$ values were estimated to be much closer ($W = 295.5, p = 0.0086$), with an estimated difference of $7.83 \times 10^{-5}$ (95% CI = $[9.27 \times 10^{-6}, 0.05]$).

### 7 Mathematical construction of shells in Main text Fig 1

The spines appearing in main text Fig 1 were formed by taking the sequence of curves, $(x_i(s), y_i(s))$, generated as output from the mechanical model, and constructing a surface by translating each curve in the transverse direction by a fixed amount, $\delta z$, i.e., creating the curves $(x_i(s), y_i(s), i\delta z)$ in $\mathbb{R}^3$, and then interpolating the curves to generate a surface. Values of $(g_1, g_2)$ for the spines were: $(0.1, 0.25)$ for Hayama (Fig 1(b)), $(0.5, 0.25)$ for Jyogashima (Fig 1(c)), and $(0.1, 0.22)$ for Tateyama (Fig 1(d)). The simulated full shell shown in Fig 1(b) was created by first constructing a smooth shell, following the procedure outlined in [9, 10], and then imposing the spine shape on top with a chosen spatial and temporal periodicity by identifying the Cartesian coordinates, $(x, y, z)$, with the local coordinates on the shell surface (in the notation of [9], we identify $x$ with the tangent direction, $\mathbf{d}_3$, $y$ with the negative of the normal direction, $\mathbf{d}_1$, and $z$ with the binormal direction, $\mathbf{d}_2$).

6. Pauli Virtanen, Ralf Gommers, Travis E. Oliphant, Matt Haberland, Tyler Reddy, David Cournapeau, Evgeni Burovski, Pearu Peterson, Warren Weckesser, Jonathan Bright, Stéfan J. van der Walt, Matthew Brett, Joshua Wilson, K. Jarrod Millman, Nikolay Mayorov, Andrew R. J. Nelson, Eric Jones, Robert Kern, Eric Larson, C J Carey, İlhan Polat, Yu Feng, Eric W. Moore, Jake VanderPlas, Denis Laxalde, Josef Perktold, Robert Cimrman, Ian Henriksen, E. A. Quintero, Charles R. Harris, Anne M. Archibald, Antônio H. Ribeiro, Fabian Pedregosa, Paul van Mulbregt, and SciPy 1.0 Contributors.

SciPy 1.0: Fundamental Algorithms for Scientific Computing in Python. *Nature Methods*, 17:261–272, 2020.

7. Wolfram Research, Inc. Mathematica, Version 13.1. Champaign, IL, 2022.
8. James D Murray. *Mathematical biology II: Spatial models and biomedical applications*, volume 3. Springer New York, 2001.
9. Derek E Moulton and Alain Goriely. Surface growth kinematics via local curve evolution. *Journal of mathematical biology*, 68(1):81–108, 2014.
10. Derek E Moulton, Alain Goriely, and Régis Chirat. Mechanics unlocks the morphogenetic puzzle of interlocking bivalved shells. *Proceedings of the National Academy of Sciences*, 117(1):43–51, 2020.

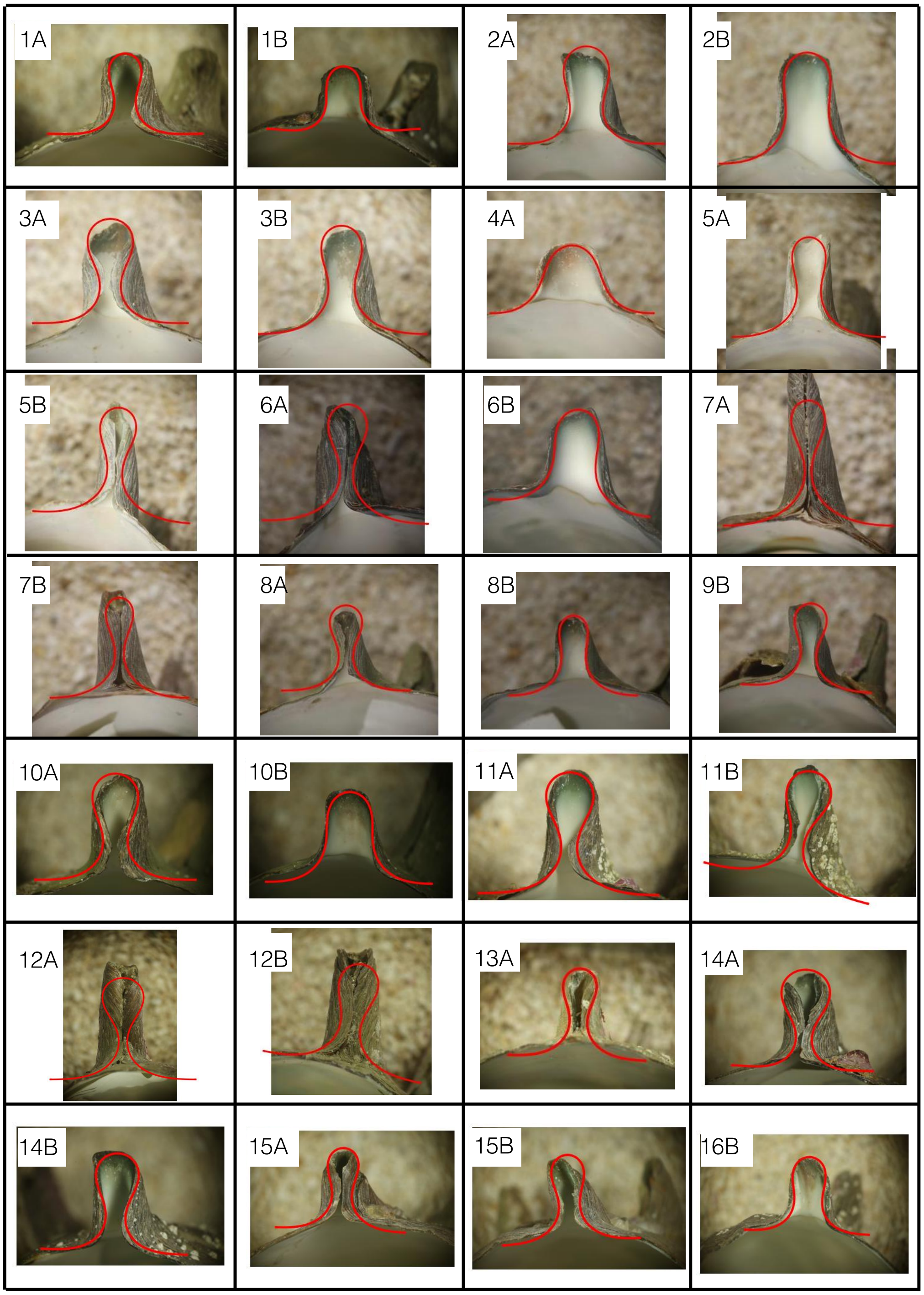


**3.** Spines analysed from Jyogashima, including a single curve from the tissue model overlaid. Extracted $(g_1, g_2)$ values are given in Table 2.

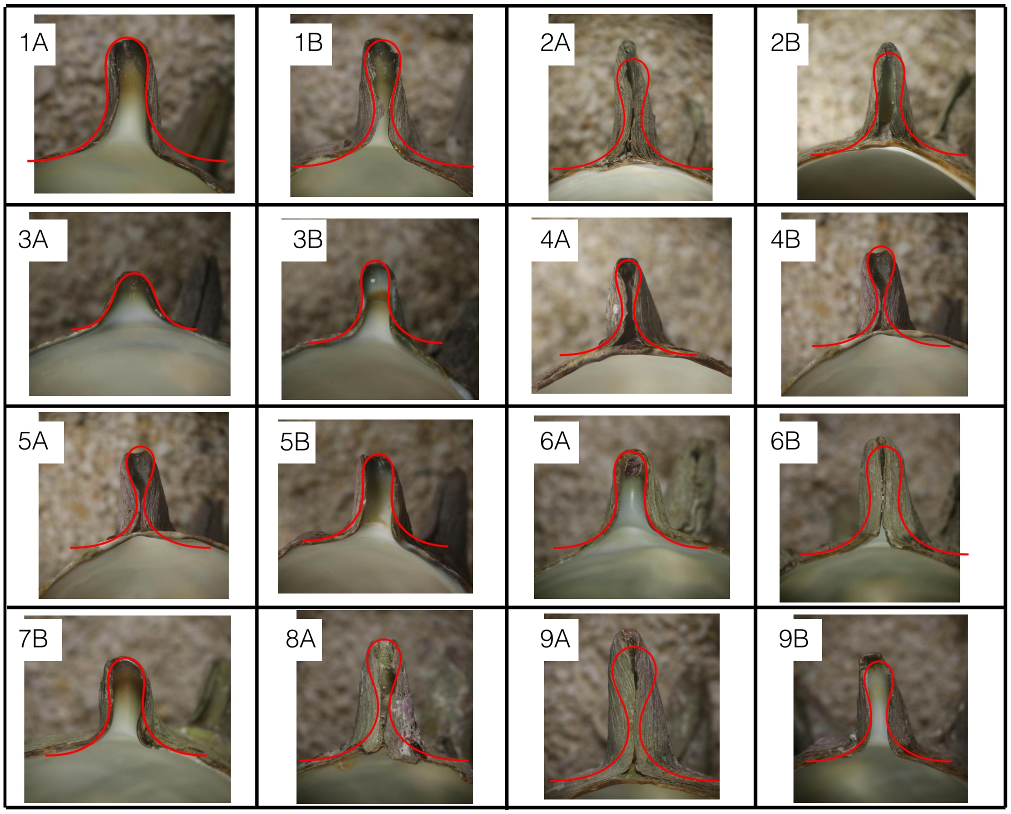


**4.** Spines analysed from Tateyama, including a single curve from the tissue model overlaid. Extracted $(g_1, g_2)$ values are given in Table 3.
